# Epidemiology of Dengue viral infection in Nepal: 2006-2019

**DOI:** 10.1101/2020.08.27.269720

**Authors:** Komal Raj Rijal, Bipin Adhikari, Bindu Ghimire, Binod Dhungel, Uttam Raj Pyakurel, Prakash Shah, Anup Bastola, Megha Raj Banjara, Basu Dev Pandey, Daniel M. Parker, Prakash Ghimire

## Abstract

**Background:** Dengue is one of the newest emerging diseases in Nepal with increasing burden and geographic spread over the last 14 years. The main objective of this study was to explore the spatio-temporal epidemiological patterns of Dengue since its first report (2006) till 2019 in Nepal.

**Methods:** This study is a retrospective analysis of dengue data available from the Epidemiological Disease Control Division (EDCD) of Government of Nepal. The data in this study cover the last 14 years (2006-2019) of reported dengue cases in Nepal. Epidemiological trend and spatio-temporal analyses were performed. Maps of reported case incidence were created using QGIS version 3.4.

**Results:** Since the first report of dengue in a foreigner in 2004, Nepal reported a total of 17,992 dengue cases in 68 districts of Nepal in 2019. The incidence was approximately five times higher in 2018 (Incidence Rate Ratio (IRR): 4.8; 95% CI: 1.5 – 15.3) and over 140 times higher in 2019 (IRR: 141.6; CI: 45.8 – 438.4). Population density was not a statistically significant predictor of case incidence. Mean elevation had a negative association with case incidence. A one standard deviation increase in elevation was associated with a 90% decrease in reported case incidence (IRR: 0.10; CI: 0.01 – 0.20). However, the association with mean elevation varied across the years. In comparison to 2016, incidence was greater at higher elevations in 2018 (IRR: 22.7; CI: 6.0 - 86.1) and 2019 (IRR: 9.6; CI: 2.6 - 36.1).

**Conclusion:** There is a high risk of dengue outbreak in the Terai region with increasing spread towards the mid-mountains and beyond as seen over the last 14 years. Urgent measures are required to increase the availability of diagnostics and resources to mitigate future dengue epidemics. Findings from this study can inform the spatio-temporal distribution of dengue and can help in resource allocation and priority setting for future epidemic.

**Author summary:** Dengue in humans is caused by four different serotypes (DENV-1, DENV-2, DENV-3 & DENV-4). Globally it is the most pervasive vector borne diseases with increasing number of cases in recent years. Dengue is one of the youngest emerging diseases in Nepal with increasing cases and spread from the tropical lowland to the highland (hilly) regions. We conducted a spatio-temporal analysis of national data to consolidate the information using QGIS to measure the dengue incidence at district levels of Nepal. Spatio-temporal analysis exploring the incidence and distribution of dengue cases aids in identification of high-risk areas which can ultimately enable national dengue programme to mobilize and allocate resources for the control and treatment. This study shows, the persistent high risk of dengue outbreak in lowland Terai region with annual rise in the risk of spread towards the mid-mountains and beyond. Urgent measures are required to increase the diagnostics and resources to mitigate the epidemic burden of dengue in Terai and peripheral regions.

## Background

Nepal has seen the outbreak of several emerging and re-emerging diseases in recent years, including dengue fever, rickettsial fevers, and other vector borne diseases [1]. The emergence of these diseases has been attributed to ecological changes, climate change, dispersion of mosquito vectors [2] and human population dynamics [1]. Nepal has three major ecological zones: the tropical Terai region, a subtropical and temperate mid-hill region, and the subalpine to alpine Himalayan region [3, 4].

Dengue fever, malaria, and Japanese encephalitis are among the most common vector borne diseases (VBDs) in low and middle income countries (LMICs) [5]. The endemicity and overall burden of VBDs in LMICs is strongly related to infrastructural weaknesses, including poor water systems, sanitation, and hygiene; and the health system to respond [6]. Recent outbreaks of dengue fever in Nepal in 2019 have alarmed the public health authorities with unprecedented spread, morbidity and mortality.

Dengue is a viral infection transmitted by female *Aedes aegypti* and *Aedes albopictus* mosquitoes [7]. The causative agent, dengue virus (DENV) belongs to the genus *Flavivirus* of Flaviviridae family of single-stranded RNA virus [8, 9]. In humans, DENV has four main serotypes: DENV-1, DENV-2, DENV-3 and DENV-4 [10]. Infection with any one of these serotypes likely confers lifelong immunity to that specific serotype [11]. Infection by a new serotype may result in severe disease [12]. Most dengue infections (up to 60%) are self-limiting [13], and are characterized by acute fever, frontal headache, vomiting, myalgia, joint pain, and macular skin rash [14]. However, some patients may develop life-threatening conditions such as acute dengue hemorrhagic fever (DHF), Dengue shock syndrome (DSS), and (multi-) organ failure [15]. In the absence of effective vaccines and antiviral drugs, symptomatic treatment and vector control programs are currently the only viable strategies for dealing with dengue infections [16, 17]. Studies so far have suggested the timely diagnosis and clinical management with intravenous rehydration are critical to mitigate the severity of infection below 1% [18]. Transmission can be reduced through protection from blood feeding Aedes mosquitoes.

The laboratory diagnosis of dengue is supported by the clinical suspicion followed by diagnostics that include rapid diagnostic tests (RDT), Enzyme linked Immunosorbent assay (ELISA) and complete blood counts (CBC) [19]. CBC profile demonstrating leucopenia, thrombocytopenia, increased hematocrit and liver enzymes are some of the parameters that aid in clinical suspicion [19]. More specific and sensitive diagnostic tools such as viral isolation and culture, and detection of viral antigen by polymerase chain reaction (PCR) are not routinely performed in Nepal [19]. Moreover, serological tools are used even during epidemic outbreak which further limit the proper diagnosis of disease in Nepal [20].

Previous studies conducted in Nepal have explored the seroprevalence in various regions since the likely initial outbreak of dengue in Nepal in 2006. Overall seroprevalence of 10.4% (anti-DENV-Ig) was found among suspected cases of DF and DHF in south-west region of Nepal 2006 [20]. Seroprevalence studies targeting smaller geographic locations have found 7.7% in Kathmandu in 2007 [21], 29.3% in south-western Terai between 2007 and 2008 [22], 9.8% in 2009 in the same region [23], 12.2% in Kanchanpur [24], 11.8% Bharatpur and Rapti Zonal Hospital in 2011 [25, 26], and 19.3% in Chitwan and Dang in 2013 [27].

Although the Government of Nepal has developed an Early Warming and Reporting Systems (EWRS) to issue warning on potential outbreaks, the response to dengue outbreaks have not been sufficient to prevent outbreaks. In 2019 there was a massive epidemic in Nepal [28], coinciding with epidemics of dengue and other Aedes-spread diseases throughout much of the tropical world. There are several challenges for prevention and control of dengue infection in Nepal, among which robust mechanism to respond to the outbreak has been constrained by lack of updated epidemiological data. In addition, Nepal has recently entered into federal system with three tiers of government: federal, provincial and local which lack effective coordination that has been adversely impacting on the management of human resources, logistic chain management and surveillance [29]. To mitigate these challenges, the federal system has devised an integrated vector control strategy (that includes diseases such as malaria, and kalaazar), currently under preparation. Nonetheless, variation in characteristics of vectors, mechanism of disease transmission and epidemiology may remain as major challenges.

Countering these challenges are critical in designing an effective dengue control and prevention program which largely relies on effective detection of the cases, diagnosis and prevention based on the surveillance data. There are no previous studies systematically exploring the epidemiological trend and distribution of the dengue cases in a nationwide scale. In addition, the rapid spread of dengue cases from southern Terai to northern mountainous region have prompted urgency in exploring the national dengue data from the government’s repository to inform the current and future dengue control program in Nepal. The main objective of this study was to explore the spatio-temporal and epidemiological patterns of Dengue since its first report (2006) till 2019 in Nepal.

## Methods

### Study Design

This study is a retrospective analysis of reported dengue case data available from the Epidemiological Disease Control Division (EDCD), under the Ministry of Health and Population, Government of Nepal. Dengue data were extracted from EDCD record. The data presented in this study represents serological diagnosis using rapid test kit (SD Bioline dengue IgG/IgM antibody up to 2015; and after 2015, SD Bioline dengue duo (dengue NS1 Ag+ IgG/IgM), Korea, IgM ELISA was used) of dengue cases. The data in this study covers a period of 14 years (2006-2019).

### Study sites

Nepal is a landlocked nation bordering India on the South, East and West; and China on the North. Since the declaration of new constitution in 2015, Nepal is divided into seven provinces (Province-1, Province-2, Bagmati province, Gandaki province, Province-5, Karnali Province and Sudurpachim Province) and 77 districts with area of 147,516 km^2^. It occupies 0.3% of the land region in Asia and 0.03% in the world. Nepal is located between 26°22’ N to 30°27 ' N and longitude 80°4’ E to 88° 12’ E. The geology of Nepal constitutes swamp Terai at 70m from ocean level to the highest elevation: Mount Everest (8848m). Land divisions incorporate Terai, Hills and Mountains. The most recent statistics in 2011 estimated a population of 26.5 million with a development pace of 1.35 individual per annum [30]. Over half of the populace lives in the Terai district of Nepal, where vector borne diseases, malaria, dengue, Japanese encephalitis, kalazar fever are endemic.

### Data collection

In Nepal, data on dengue surveillance is collected by the health system infrastructure that includes Health Post, Primary Health Center (PHC), District Hospital, Provincial Hospital and Central Hospital. Dengue cases recorded in the health center are collected monthly and are reported to the District Health Office (DHO)/District Public Health Office (DPHO). The information subsequently is notified to the Epidemiology and Diseases Control Division (EDCD) from DHO/DPHO on a monthly basis through Health Management Information System (HMIS)-reporting mechanism. Besides HMIS, an Early Warning Reporting System (EWARS) is utilized to record hospital admitted dengue cases and dengue deaths. Population density at the district level was calculated as people per km^2^. District level population counts were derived from the 2011 Nepal Census. We calculated mean elevation for each district using elevation data from the GTOPO30 global digital elevation model (DEM).

### Data analysis

Data were first entered in Microsoft Excel 2010 and later analyzed. The trend of incidence of dengue cases and proportions of dengue cases in different province/districts in Nepal (2006-2019) were analyzed. The map of dengue cases of 2019 was created using QGIS software version 3.4 (https://qgis.org/en/site/). We used a mixed effects negative binomial regression in order to test for associations between calendar year, mean elevation at the district level, district population density. A random intercept was used for district to account for repeated observations within each district across the calendar years. The outcome variable was reported case incidence per 100,000 per year, rounded to the nearest whole number. We hypothesized that incidence was increasing at higher elevations over time and included an interaction term between calendar year and mean elevation to test our hypothesis. Both population density and mean elevation were centered on their mean values and standardized using their respective standard deviations so that a one-unit change in both values corresponds to one standard deviation change.

### Software

Maps of reported case incidence were created using QGIS version 3.4. Population density and mean elevation at the district level were both calculated in QGIS. The negative binomial regressions were done using R statistical software version 3.5.2.

### Ethics

This study obtained ethical approval from Ethical Review Board of Nepal Health Research Council (Reg no. 496/2020 P).

## Results

### Annual trend of dengue incidence in Nepal (2006-2019)

The trend of dengue (confirmed by serological test either IgM ELISA or rapid test kit) incidence over the period of 2006-2016 was analyzed (Fig 1). The trends of dengue incidence are presented below in three-year periods to divide the equal proportion of time period.

**Figure 1.**
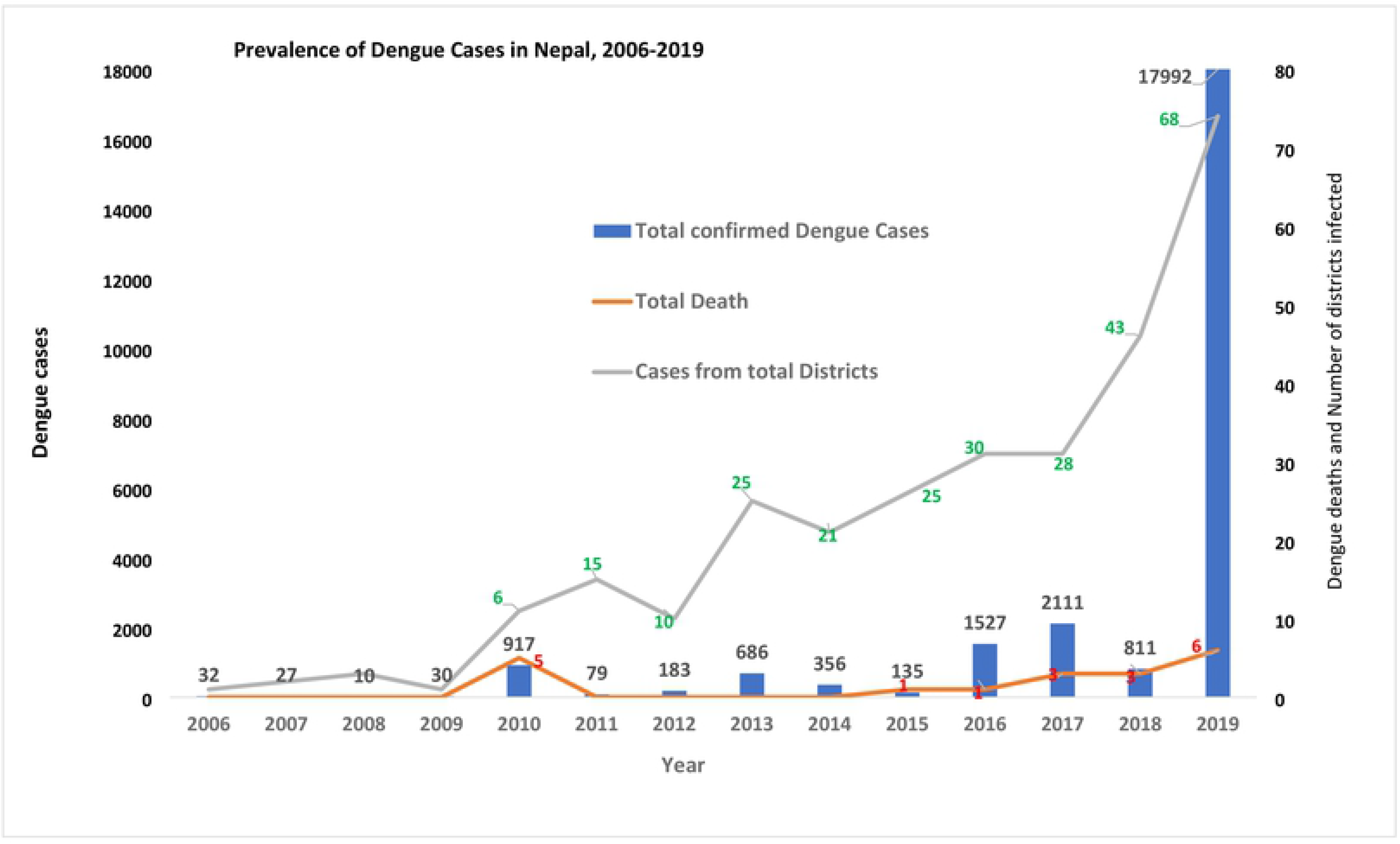
Prevalence of Dengue in Nepal:2006-2019.

### Dengue in 2006

Nepal reported its first dengue case in a Japanese foreigner, imported from India in 2004. Two years later there was an endogenous outbreak in lowland Chitwan district in 2006; with a total of 32 reported dengue cases throughout the country.

### Period between 2007 and 2010

From 2007 to 2009, the total number of reported dengue cases was low in comparison to the 2006 epidemic. In 2007, 27 dengue cases were reported from 4 districts of Terai region. However, there was slight decrease in number of dengue cases; reported 10 from three districts. In 2010 Nepal had faced major outbreak of dengue. The dengue cases rose to 917 by 2010 with 5 reported deaths (2 from Chitwan district, 1 from Nawalparasi and 2 from Rupandehi district) from six districts of Nepal (Figure1).

### Period between 2011 and 2013

In 2011, the number of dengue cases (79 cases) were very low in comparison to 2010. However, there was an expansion in its distribution, cases were reported from 15 districts of Nepal (only 6 districts in 2010 epidemic). There was another dengue epidemic in 2013 and total of 686 cases were reported from 25 districts of Nepal. There was no report of death due to dengue between 2011 and 2013.

### Period between 2014 and 2016

In 2014, a total of 356 dengue cases were reported from 21 districts of Nepal. Out of the 356 cases, 50.8% (181/356) were from Bagmati province, 35.9% (128/356) from province-2, 7.3% (26/356) from Sudurpachim province, 3.6% (13/356) from Province-5 and 2.2% (8/356) from province-1. There were no dengue cases reported from Karnali province and Gandaki province in 2014 (Table 1). In 2015, the total reported dengue cases were only 135 throughout the country with majority from Bagmati province (76 cases out of 135 cases) and 1 death from Dang district. In 2016, there was another dengue epidemic in Nepal and a total of 1527 dengue cases were reported from 30 districts; comprising all seven provinces. Only one dengue death was reported from Chitwan district in 2016 epidemic. Province wise dengue cases distribution in 2016 showed 51.2% (781/1527) of cases from Bagmati province, 27.4% (418/1527) from province −1, 15.8% (242/1527) from province-5, 2.8% (43/1527) from province-2, 1.5% (23/1527) from Karnali Province, 1.1% (17/1527) from Sudurpachim Province; and only 3 dengue cases were reported from Gandaki province (Table 1). Among 1527 cases, 44.8% (687/1527) were from Chitwan district and 26.5% (405/1527) were from Jhapa district in 2016. (Supplementary Table 1).

### Period between 2017 and 2019

In 2017, a total of 2111 dengue cases were reported from 28 districts of Nepal. The number of cases rose by 38% (1527 in 2016 versus 2111 in 2017) in comparison to 2016. Out of 2111 cases, 40.4% (853/2111) were from province-5, 28.8% (609/2111) from province-2, 25.7% (543/2111) from province-1, and 4.5% (95/2111) from Bagmati province. There were three dengue deaths reported each from Palpa, Chitwan and Makawanpur districts in 2017. In 2018, the total reported dengue cases were only 811 throughout the country with majority from Gandaki province (568 cases out of 811 cases). There were 3 deaths reported in Rupandehi (2 cases) and Makawanpur district (1 case) from dengue in 2018 with distribution expanded to 43 districts of Nepal.

In 2019, there was another huge dengue epidemic in Nepal with a total of 17,992 dengue cases with expansion to 68 districts of Nepal; comprising all seven provinces. There were six dengue deaths reported from 5 districts of Nepal (2 deaths in Chitwan, and 1 each death in Sunsari, Sindhupalanchock, Kathmandu and Doti) in 2019 epidemic. On province wise dengue cases distribution in 2019, 40.5% (7276/17992) were from Bagmati province, 24.4% (4379/17992) from province −1, 19% (3421/17992) from Gandaki province, 13.4% (2414/17992) from province-5, 1.5% (276/17992) from province −2, 0.8% (152/17992) from Sudurpachim province and very low (0.4%; 74/17992) from Karnali province (Figure 2) (Table 1). On district wise distribution, Sunsari district comprised 19% (3431/17992) followed by Chitwan (18.9%; 3402/17992), Kaski (15.7%; 2824/17992), Kathmandu (8.8%; 1589/17992), Lalitpur (3.3%; 596/17992) and Jhapa (2.9%; 525/17992) (Supplementary Table 1).

**Figure 2:**
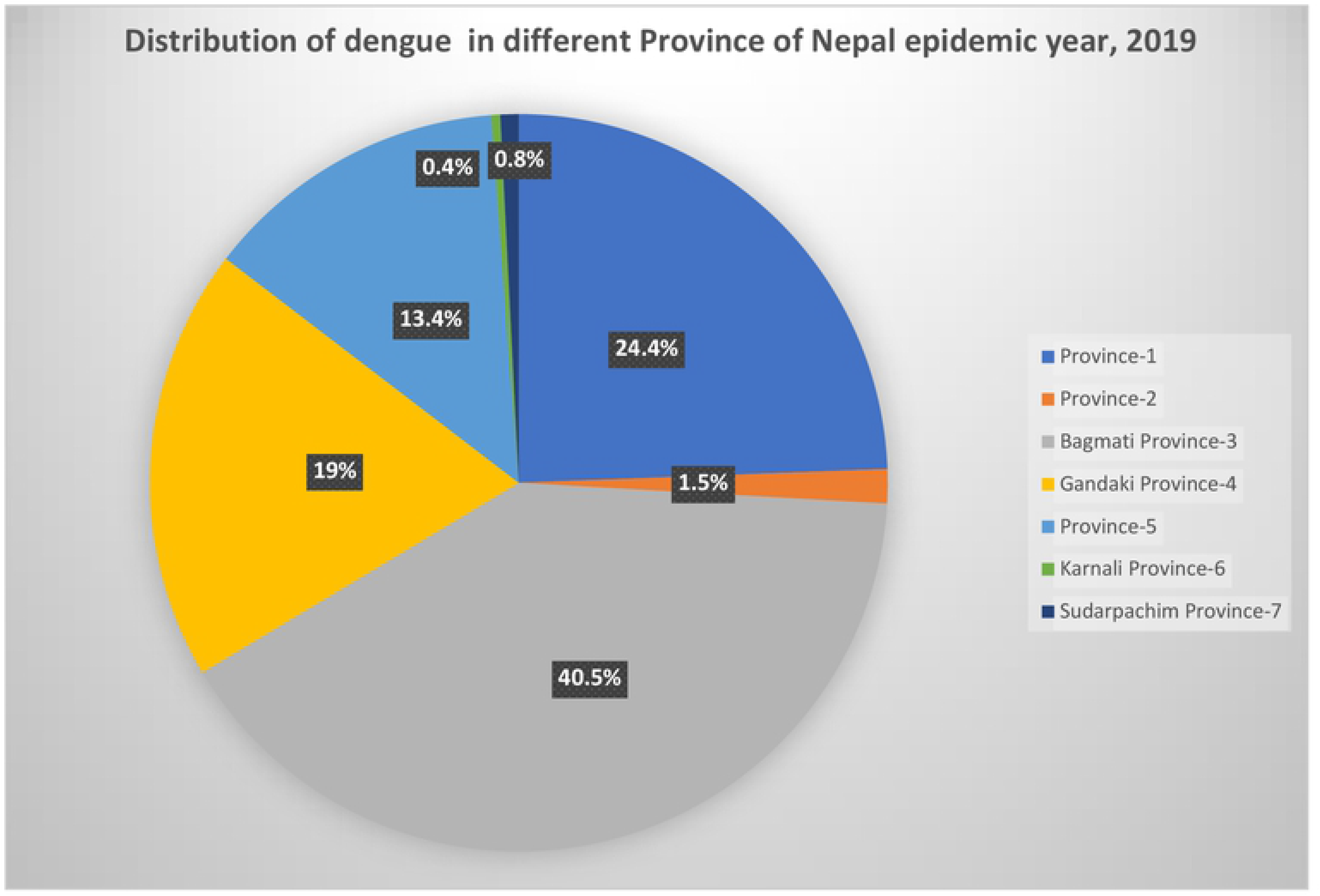
Percentage distribution of dengue cases in different provinces of Nepal in 2019.

### Choropleth maps of reported dengue fever case incidence at the district level (2016 – 2019) and dengue incidence over time

Choropleth maps of reported dengue fever case incidence at the district level (2016 – 2019) were generated, with case incidence presented as the number of cases per 100,000 people for each year (Figure 3).

**Figure 3:**
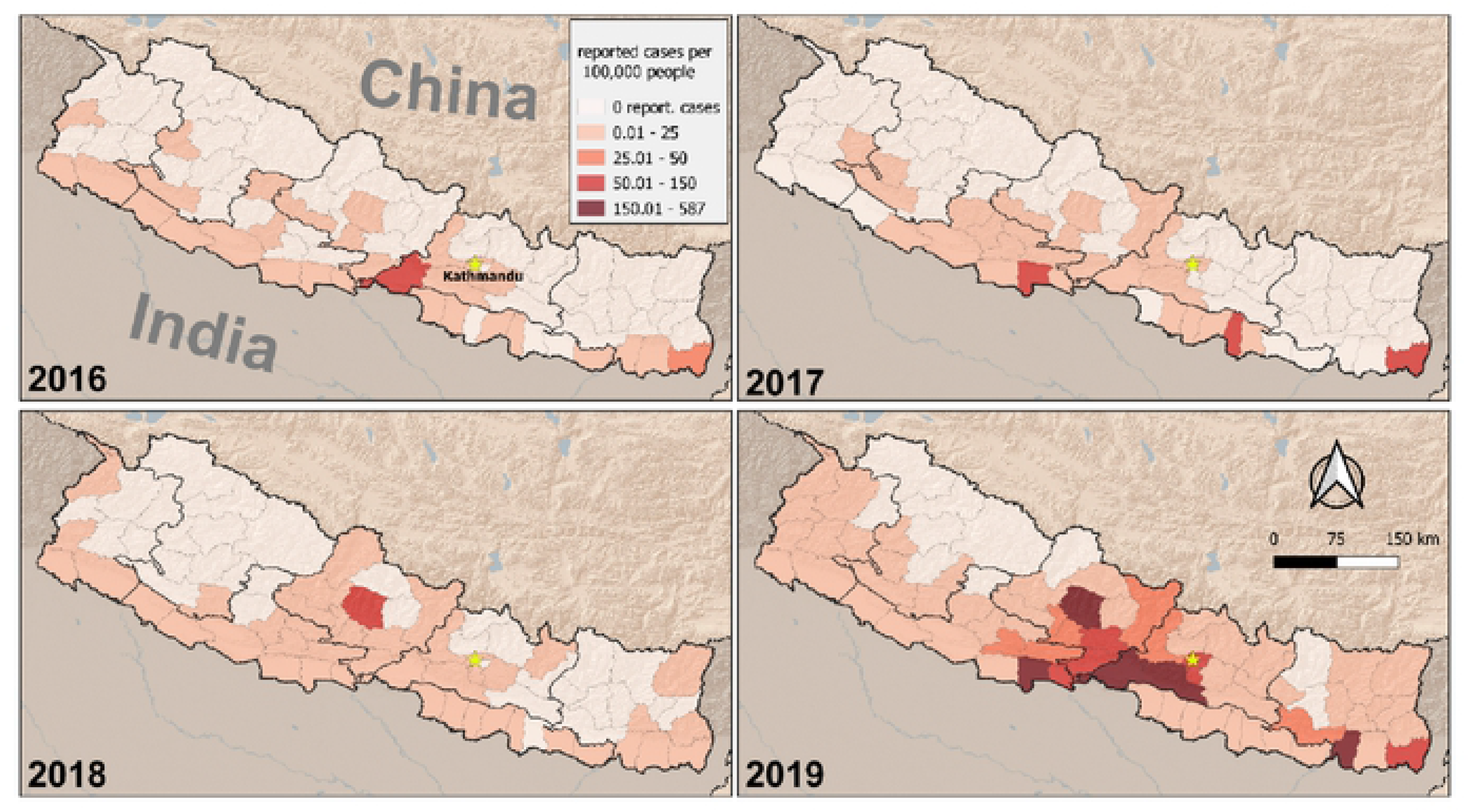
Choropleth maps of reported dengue fever case incidence at the district level (2016 - 2019). case incidence is presented as number of cases per 100,000 people for each year.

Reported case incidence was much higher in 2018 and 2019, using 2016 as a comparison (Table 3). The incidence was approximately five times higher in 2018 (Incidence Rate Ratio (IRR): 4.8; 95% CI: 1.5 – 15.3) and over 140 times higher in 2019 (IRR: 141.6; CI: 45.8 – 438.4). Population density was not a statistically significant predictor of case incidence. Mean elevation had a negative association with case incidence (Table 2). A one standard deviation increase in elevation was associated with a 90% decrease in reported case incidence (IRR: 0.10; CI: 0.01 – 0.20). However, the association with mean elevation varied across the years, as is evident from the interaction effect in our model. In comparison to 2016, incidence was greater at higher elevations in 2018 (IRR: 22.7; CI: 6.0 - 86.1) and 2019 (IRR: 9.6; CI: 2.6 - 36.1).

## Discussion

### Overview of findings

Since the first report of dengue in 2004 [31], Nepal has steadily experienced the rise and expansion of cases, with 17992 cases in 2018/2019 and has spread in majority of the districts (68/77). The four-year choropleth maps (2016-2019) of reported dengue fever case incidence at the district level showed dengue incidence was five times higher in 2018 and over 140 times higher in 2019. Such a steady rise and nationwide distribution of dengue makes the disease a national priority with urgent implications for control and prevention.

### Trends of Dengue cases in Nepal

The overall trend of dengue incidence and its distribution show a rising trend with outbreaks in 2010, 2013, 2016 and 2019 in Nepal. In just last six years since the first imported case of dengue in 2004, Nepal has become an endemic country for dengue. Following the outbreak of DF/DHF in India in 2006, a minor outbreak was confirmed in the same year from nine districts of Nepal [20, 22] with 32 cases but no fatalities. In 2006 the diseases was reported from nine districts from lowland tropical Terai and hilly districts such as Kathmandu were spared from the outbreak [20]. The primary vector of dengue transmission, *Ae. aegypti,* was reported from only five districts implying a possibility of high importation of cases [19].

Dengue remained almost latent during the period of 2007 and 2009, until the massive outbreak of 2010 with 917 cases and distribution into six districts. A study conducted in southern Terai during this period showed a high prevalence (29.3%) of anti-DENV IgM [22] which however was countered by a subsequent study that showed lower prevalence of 9.8% in 2009 [23]. In the same year, with an extensive cross-sectional study covering southern Terai showed an overall seroprevalence of 12.1% with high proportion in Kanchanpur bordering India [24]. These studies showed the high vulnerability and impending epidemic outbreak in the Terai region [24]. Around 80% of the total confirmed cases were reported from Terai region that showed all serotypes with entomological evidence of both vectors: *Ae. aegypti* and *Ae. albopictus* [20].

The outbreak of 2016 was the result of re-emergence of DENV-1 that recorded 1527 cases, with distribution in 30 districts of Nepal. Two terai districts: Chitwan and Jhapa accounted almost 72% (1092/1527) of the total cases registered. Both of these districts have a tropical climate, and border with India which can explain in part the high incidence of dengue cases [32]. Following the first report of dengue in highland region in 2010, 3.1% of the total cases, with 0.4% from Kathmandu alone were reported by the end of 2016. National Public Health Laboratory (NPHL) Kathmandu reported 16.9% (45 out of 266) of patients showing anti dengue IgM antibodies in serum [33].

Triennial peak and expansion in distribution of the dengue epidemic of 2010, 2013, 2016 and 2019 are in line with the previous reports from Brazil [34] and Cuba [35]. In the subsequent outbreaks, serious complications associated with the dengue infection were not observed as expected and could be due to prevalence of newer serotype DENV-2 in the 2013 outbreak [36]. This could be due to the low virulence of newer strain, or the cross-immunity developed due to each endogenous infection. Also, the higher mortalities and morbidities are associated with secondary infection with cross-infections with another serotype. Antibody dependent enhancement has been shown to be causing severe form of dengue, also known as secondary explosion as is observed in India, Bangladesh; [37] Vietnam, Singapore and Senegal [38]. The same mechanism may have a role in the outbreak that occurred in 2016 and 2019 in Nepal.

### Geospatial distribution of dengue cases

In 2010, DENV-1 was the prominent serotype for the epidemic. However, the outbreak of 2013 was solely caused by the DENV-2 [36, 39]. This indicated the prevalence of all serotypes with endemicity of DENV as silent threat all over the country. Three lowland Terai districts--Chitwan, Jhapa, and Parsa were again worst hit districts, together constituting the 85% of total cases in the country. Over the years, rising prevalence of dengue in Kathmandu has countered the presumption that Kathmandu was climatically unsuitable for dengue vector. Increased urbanization, industrialization together with the climate change may have contributed a conducive ambient environment for *Aedes* vector mosquitoes [40–42].

The outbreak of 2016 showed both increase in the number of cases and the distribution of disease to newer temperate zones within Nepal. For instance, temperate hilly zones of Gandaki province began to report cases in 2015 while the outbreak in subsequent year affected Karnali province located in upper hilly region. Also, the outbreak of 2016 marked the geographic expansion of dengue infection in all seven provinces. The emergence and re-emergence of DENV serotypes intermittently in varying manifestations implies the possible burst of severe form of the dengue related illnesses. Similar mechanism and patterns of DENV infection with multiple virus clades were observed in Indonesia [43] and Brazil [44] while circulation of DENV-1 in the same period (2014-2016) was also observed in China [45] and other South Asian countries: India [46], Bangladesh [47], Pakistan [48] and Sri Lanka [49]. In our study, dengue case incidence showed five times higher incidence in 2018 and over 140 times in 2019 in comparison to 2016 (Table 3). The findings of our study are in line with studies reported from Thailand [50] and Bangladesh [51].

There was a steady rise in number of cases and its distribution between the period of 2017 and 2019. The outbreak is remarkable for its spatial and temporal shift in addition to the role of two serotypes (DENV-1 & DENV-2) [1]. Kathmandu saw the repeated outbreak of dengue and since then experts fear the imminent outbreak of dengue in future. Although the vectors are capable of flying only 500m in their lifetime [52, 53], a number of underlying factors such as urbanization, trade and transit from dengue-infested regions and climate change are favoring their spread and potency. Specifically, changes in temperature and rainfall in upland hilly regions and relative humidity in lowland plains are established as contributing factors for rise and distribution of vectors [54].

### Impact of climate change and ecology

The primary vectors: *Aedes aegypti* and *Aedes albopictus—*depend upon temperature and precipitation for their growth, survival and feeding behavior [55] and also affects vector-human transmission cycle [56]. Increasing temperature in the region can provide a favorable environment for dengue vectors and its transmission. The latest dengue outbreak of 2019 may have been flared up by unexpected early rains which may have accelerated the outbreak as early as on May 13, 2019 from Sunsari district [28]. Similarly, annual monsoon season of each year in the country makes ambient room for mosquitoes by its high humidity while the post-monsoon period favors their breeding and transmission by high rainfall and heavy flooding [1]. Some prevailing findings have suggested the existence of dengue vectors (*Aedes aegypti* and *Aedes albopictus*) from the tropical lowland to the highland Dhunche, Rasuwa (2100m elevation) district in Nepal [57]. This geographical expansion of Dengue is the result of capability of its vector to expand its geographical range of adaptability solely due to climate change caused by global warming [54].

### Implications for National Dengue control program

The government of Nepal has released the national guidelines for the prevention, control, and management of dengue in the country which has focused the vector-control strategies as the best policy to curb the potential epidemics. Despite the guideline, the rising trend of dengue cases with expansion in distribution in almost all the districts of Nepal poses significant challenges. Of the challenges, Nepal can plan through the historical account of dengue epidemiology, rising trend and its spread in the districts. Specifically, the visualization of dengue cases among the districts can also help in categorizing and prioritizing the districts based on the epidemiological burden identified in this study. Also, the dengue control and prevention program can incorporate spatially focused strategies to ensure the preventive measures such as distributing mosquito repellant nets, clearing of puddles (or water collection around the households), bushy areas, and fumigation. Targeted programs with allocation of resources for treatment and prevention can be planned based on the spatio-temporal distribution of the cases visualized in this study. In addition, integrated vector control programs may benefit from the comprehensive data of dengue cases and its distribution for resource allocation.

### Strengths and limitations

This is the first study consolidating the national dengue data since the first report of dengue in Nepal till date. In addition to integrating all the data through epidemiological analysis, this study reveals the trends, and distribution of the cases which can inform the dengue control and prevention program of Nepal for instance in allocating the resources and prioritizing the preventive measures such as destructing breeding sites in the villages. This study has several limitations. Due to resemblance with other symptoms of tropical diseases, dengue cases may have been undetected and overlooked, posing a challenge on reporting [53]. The study relied on the retrospective data from government’s EDCD which may have missed the private sector data, thus may not reflect the true epidemiology of dengue in Nepal. During the outbreaks due to logistic shortages, reporting of the cases were not uniformly confirmed by RDTs, sometimes were based on the clinical diagnosis.

## Conclusions

Nepal is at major risk of burgeoning burden of dengue. Our study shows, the high risk of dengue outbreak especially in the Terai region with increasing spread towards the mid-mountains and beyond. Chikungunya and Zika viral infections are both spread by the same vectors, *Ae. aegypti* and *Ae. albopictus*. Therefore, there will be chance of spread of these infections since dengue has become endemic. Urgent measures are required to increase the diagnostics and resources to mitigate the epidemic burden of dengue in Terai and peripheral regions. In addition, limited research and surveillance in dengue has been a major hindrance to prevention and effective management. Findings from this study can inform the national dengue control and prevention program in resource allocation and priority setting with implications for future epidemic.

## Consent for publication

Not applicable

## Availability of data and materials

All data pertaining to this study are within the manuscript.

## Authors’ contribution

KRR, BA and PG developed the concept of research work. BG, BD and KRR conceived and designed the study, collected the retro specific data, carried out research works. KRR, BA, MRB, DMP analyzed the data. URP, PS, AB, MRB, BDP and PG provided oversight for the project. KRR, BA prepared the initial draft of the manuscript. KRR, BA, DMP, BDP and PG revised the subsequent versions of the manuscript. All authors read and approved the final version of manuscript.

## Acknowledgements

The authors would like thank Epidemiology and Diseases Control Division (EDCD), Department of Health Service, Ministry of Health and Population, Teku, Kathmandu.

